# Rejection in early life is associated with high and long-lasting stress and higher incidence of infections in owl monkeys

**DOI:** 10.1101/2020.10.17.343921

**Authors:** Mahdiyah Osman, Aylin Olkun, Angela M. Maldonado, Jordi Lopez-Tremoleda, Nofre Sanchez-Perea, Ursula M. Paredes

## Abstract

1.

**Background:** The mothers of owl monkeys sometimes reject their new-borns. When this occurs in captive colonies, the rejected infants are manually raised by veterinarians, allowing them to survive. However, maternal rejection can induce chronic stress, which in turn is associated with infectious diseases. Rescued, rejected owl monkeys might experience high rates of stress and infections which go undetected.

**Methodology:** To test this hypothesis, we evaluated the connection between maternal rejection, stress and infections in owl monkeys (*Aotus nancymaae*) from the IVITA Center for Conservation and Reproduction of Primates (UNMSM, Peru). Specifically, we compared the stress rates and frequency of infection treatment in juveniles (19-24 months) rejected in the first month of life and controls. To assess stress, we compared cortisol levels in hair using a competitive ELISA and recorded behaviours using cameras. We analysed past medical treatments and medication to compare incidence of infection treatment in subjects. We then studied the correlation between the frequency of infection treatments and cortisol using a linear regression.

**Results:** Rejected owl monkeys showed significantly higher cortisol levels (p=0.0123), a higher incidence of stereotypical behaviour and overeating (pacing p=0.0159; head twirling p=0.0476, eating p=0.0238) compared to controls. Rejected owl monkeys also received significantly more treatment for infections than controls per month lived (p=0.0009). Moreover, infection rates observed in this population were positively although weakly associated with concentration of cortisol in hair (R^2^=0.307, p=0.0075).

**Conclusion:** Maternal rejection in the first month of life is associated with high and long-lasting stress levels and infections in the IVITA owl monkey colony. IVITA owl monkeys will be a useful model for studying the long-term effects of early life stress at the population level.

## 2. INTRODUCTION

Owl monkeys (*Aotus s*pp.) are a genus of New World primates that are bred in captivity throughout the world as biomedical models for human infectious diseases. Breeding colonies of *Aotus* spp. suffer from high levels of maternal rejection of new-borns (Gozalo & Montoya, 1990; Sanchez et al., 2006). For primates, exposure to stressful experiences in early life has serious consequences. In nature, rejection often results in infants’ death (Mastrepieri & Carroll 2000), while in captivity, primates are hand-raised by veterinarians to facilitate their survival. Maternal rejection is associated with high stress and this in turn is related to poor health (Sanchez, 2006). Therefore, rescued, rejected owl monkeys could suffer from undetected stress, which if unchecked, could affect the well-being of these captive monkeys and introduce undesirable variations in biomedical experiments.

The number of maternal rejections in the IVITA colony of the owl monkey (*Aotus nancymaae*) has decreased considerably since its establishment, today being a successful breeding program, while maintaining international standards of animal welfare (Montoya, 2003). However, rare events still transpire. The consequences of rejection on the health and disease for these primates are unknown. Separation from the mother at an early age, whether this be experimental or spontaneous, is one of the strongest emotional stressors for a young primate. The stress caused by this experience can cause long-term deterioration in the activity of the Hypothalamic Pituitary Adrenal (HPA) axis, the system that regulates the physiological response to stress (Sanchez, 2006). While the activation of the stress response has evolved as a survival mechanism to cope with threats, chronic hyperactivity of this system is associated with aberrant behaviours and disease (McEwen, 2000). Consequently, primates separated from their mothers are more anxious. For example, maternal separation experiments on common marmosets (*Callithrix jacchus*) show a reduction in cortisol production, greater reactivity to stress and modified gene expression in neural networks involved in learning and emotion processing which persist until adolescence (Dettling et al., 2002a, Law et al., 2009). Young squirrel monkeys (*Saimiri sciureus*) temporarily separated from their mothers also show elevated cortisol levels, which persists over time (Levine et al., 1993).

Chronic stress, as the caused by experiencing maternal separation, is associated with increased risk of infections (Cohen et al., 2007). One of the causes proposed for this phenomenon is glucocorticoid resistance induced by elevated and chronic release of cortisol which impairs immune and inflammatory responses (Cohen et al., 2012). Few studies have proven the link between maternal deprivation and risk of infection in primate populations, however, retrospective studies in captive bred primates suggest a consistent effect. For example, Conti and colleagues (2012) conducted a retrospective study of health outcomes in macaques (*Macaca mulatta*) separated from their mothers, and these primates were shown to have a higher incidence of treatment for gastrointestinal infections. Similarly, Clay et al., (2015) described a lifetime elevated risk for developing upper respiratory infections in nursery reared chimpanzees (*Pan troglodytes*).

Spontaneous maternal rejection within captive breeding colonies has been less studied, perhaps because it is believed to rarely occur given adequate housing and enrichment (Nicolson, 1991) This perception is biased since much of the literature on this topic is based on Old World monkeys and apes. As for captive New World monkeys, rejections and other forms of aberrant paternal behaviour are more common. For example, Debyser (1995) reviewed causes of juvenile mortality in New World monkeys and showed that aberrant parental behaviour was one of the two leading causes of primate death before weaning age. Johnson et al., (1991) similarly reported that captive tamarins (*Saguinus oedipus*) reject their newborn infants in high numbers. It is unknown whether the higher levels of rejection reflect a particular sensitivity of New World monkeys to stressors or different maternal care strategies in this taxon. Given the numerous breeding colonies of New World monkeys, the need to investigate this phenomenon remains urgent.

In the IVITA owl monkey colony, rejection was previously associated with higher mortality amongst juveniles (Gozalo & Montoya, 1990; Sanchez et al., 2006). Whereas hand rearing has eliminated mortality of rejected owl monkeys (Personal communication Sanchez, N), whether these individuals suffer from more infections which can threaten survival of juveniles is unknown. Given the presented evidence, we hypothesized that rejected and hand reared owl monkeys suffer from chronic stress and are more likely to suffer from stress-related infections. The objective of our study was to test whether there is an association between maternal rejection, elevated stress and infections history in juvenile owl monkeys from the IVITA colony. To accomplish this task, we compared stress biomarkers (physiological and behavioural: cortisol and stereotypies respectively) and records of administration of medications given for infections in rejected juveniles (in the first month of life) and age matched controls. Our analysis demonstrates that rejected owl monkeys have elevated cortisol levels and exhibit behavioural signs of distress, requiring more treatments for bacterial infections even two years after rejection. Our report supports that preventing maternal rejection and / or understanding its effects would be key to improving and refining health management and prevention of disease in owl monkey colonies.

## 3. MATERIALS AND METHODS

### Study setting

We studied the *Aotus nancymaae* colony housed in the facilities of the Center for the Reproduction and Conservation of Non-Human Primates (CRCP) of the Veterinary Institute of Tropical and Altitude Research (IVITA) of the Universidad Nacional Mayor de San Marcos (UNMSM). This is located in the city of Iquitos, Department of Loreto, the Amazon region of Peru. IVITA’s owl monkey colony is made up of 1,500 monkeys, which originated in the tropical forests that surround the city of Iquitos. The monkeys are housed in large buildings open to the environment with metal mesh doors that allows exposure to the natural habitat bioperiod throughout the year. This species is monogamous, and in the wild forms family groups made up of breeding pairs and descendants (Aquino & Encarnacion; 1986). IVITA enclosures contain a single breeding pair or breeding pair plus offspring. The enclosures have a larger vertical surface (2m3, width = 1m, depth = 1m height = 2m) as recommended for housing arboreal New World monkeys (National Research Council, 1998). The enclosures contain a family nest located in the upper left corner and 3 perching branches for vertical and horizontal movement.

### Family and rearing environment

In IVITA the monkeys live in family units formed by a mother and father parental unit and young until they reach 7 months which emulates observed family organization in nature (Aquino & Encarnacion 1986). A couple of days after birth, infants begin to be carried by their fathers who are the primary care givers in this species, other than breastfeeding (N. Sanchez, personal communication). At 7 months, the juveniles are fully weaned from breast milk and move independently. At this age, juveniles are moved to a new home and are paired with members of the opposite sex.

### Maternal rejection

Mothers sometimes reject their young shortly after giving birth (within the first four weeks of life). This is characterised by mothers that block access to the nipple for suckling, ignore calls for help, and sometimes bite the tail and toes in order to persuade the young to stop searching for the nipple. Whereas this form of rejection is considered part of the normal weaning process for *Aotus* spp. (Dixon, 1982), it can escalate to abuse and be lethal (Gozalo & Montoya, 1990; Sanchez et al., 2006). If this occurs, rejected infants are transferred to incubators and veterinarians raise them by hand. The vast majority of individuals transferred to the incubators are successfully bred and re-join the breeding colony at 7 months. For the rejected group, food consists of human infant formula (Nan-Nestlé, USA) until 4 months of age after which formula milk is combined with IVITA’s own owl monkey biscuits (composed of: wheat flour, soy protein, peanuts, vitamins, minerals and sugar) and supplemented with seasonal fruit.

### Ethical statement

All procedures for handling monkeys, sampling and behaviour observations were approved by UNMSM’s ethics committee. The collection of samples for this study was completed following SERFOR regulations, Peru’s national forest and wildlife authority. Authorization for sampling was granted to Angela Maldonado and Ursula M. Paredes (SERFOR RDG0050-2018-SERFOR-DGGSPFFS). The samples were processed at the Dr. José Pérez Zúñiga Ecology and Conservation Laboratory of the Cayetano Heredia University, Lima, Peru.

### Animal samples

To draw blood samples and cut hair for cortisol testing along with regular IVITA veterinary wellness check-ups, the animals were anesthetized with ketamine hydrochloride (10-20 mg/kg) intramuscularly following established methods (Sánchez et al., 2018). The criteria for the inclusion of samples for the rejected group were: juveniles of both sexes, 19-24 months old, rejected within the first month of life. The criteria for the controls were: 19-24 month-old juveniles of both sexes born 0-1 days around the birth of rejected juveniles. A blind analysis of samples was carried out at Queen Mary University of London, UK.

### Cortisol measurements

To measure cortisol levels in hair, 1.5 cm strands were cut from the tip of the tail. The hair was stored in the refrigerator at 4°C until processing time. Hair was cut into small sections with scissors and then weighed in a microbalance. To isolate and measure cortisol, an Isopropanol precipitation protocol was used, followed by a competitive ELISA salivary cortisol (Salimetrics, USA) validated for hair (Meyer et al., 2014). Immunoassays were measured as colour change on a plate reader following the manufacturer’s instructions. The protocol included a standard curve. The raw values were converted to concentrations using a reciprocal absorbance method, using the slope value obtained from the standard curve.

### Behaviour recordings

*Aotus nancymaae* is a nocturnal primate whose activity peaks between 5 pm and 6 am (Aquino & Encarnacion 1986). Therefore, we recorded nocturnal behaviours with motion-sensitive infrared cameras located in 10 enclosures, 5 containing abandoned, hand-reared individuals and 5 enclosures with controls. As individuals were not differentiated, the behavioural recordings were based on households rather than individuals. The motion-sensitive cameras (Trail Wildlife Camera Trap with Infrared Night Vision, APEMAN 12MP 1080P) were placed 1 m from the floor and 1.5m from the front wall, offering a complete view of the enclosures. Cameras recorded during 5 consecutive nights, with recordings made between 5:30 pm and 5:30 am, (total recorded time for rejected=17.69 hours, controls = 9.11 hours), spanning 60 hours. An ethogram was developed to capture the prevalence of behaviours such as feeding, resting, playing, grooming, and stereotypy: head twirls and pacing from one side to the other. The behaviours counted were scored by two independent observers.

### Medical histories

IVITA’s veterinary health records include endo and ectoparasite screenings, body condition, injuries and medication that provided information to determine the age at which individuals were rejected and drugs prescribed. All drugs to treat bacterial, fungal and parasitic infections (e.g. amoxicillin, sulfonamide, cephalosporin, trimethopine, metradinazole, ivermectin) were quantified for this study. To compare drug administration in individuals of different ages, the number of interventions was normalized per number of months lived to the date of collection of biological samples.

### Statistical analysis

We performed the non-parametric Wech’s t-test to compare statistical differences between the median values of cortisol levels, the incidence of behaviours and the frequency of medication of rejected and controls. The minimum value at which null hypotheses were rejected was set at 0.05.

## 4. RESULTS

Our study is the first to show that owl monkeys rejected by their mothers in the first month of life are more stressed and receive more medication for infections than controls. These effects are long-lasting, and are detected in juveniles almost up to 2 years post-rejection. Our study suggests a weak correlation between cortisol levels and infection treatment for owl monkeys.

Cortisol concentration per gram of hair was compared using an ELISA immunoassay between rejected and controls (Figure 1.A). Our analysis identified a significant variation in concentration in both groups (rejected = 41.47 ± 3.29; controls = 30.97 ± 14.27). However, an unpaired Welch’s t*-*test demonstrated that there were statistically significant differences between the mean hair cortisol value of rejected and control monkeys (two-tailed, t = 2.838, df = 15.21, p = 0.0123).

**Figure 1.**
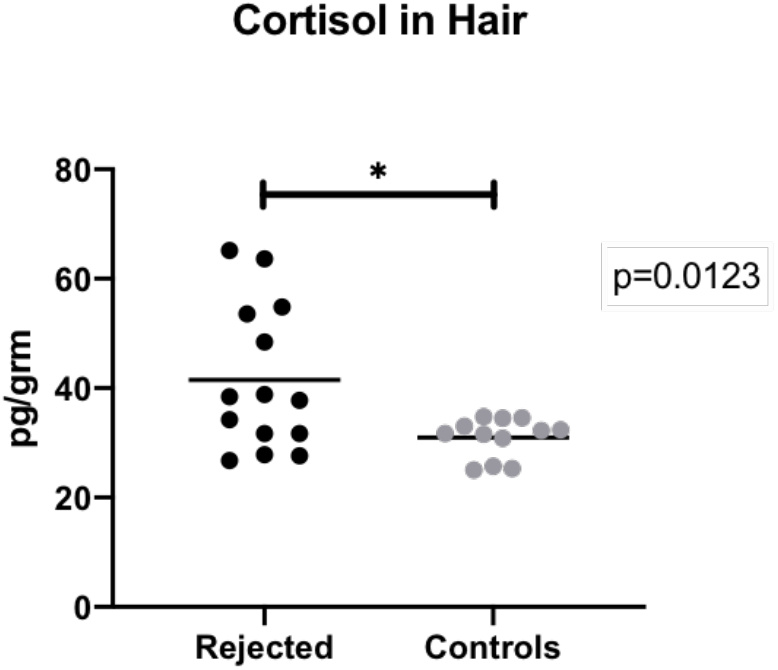
Concentration of cortisol in the hair of owl monkeys IVITA. Rejected Juveniles (black circles) have higher cortisol concentration than controls (grey circles), Probability *p* derived from a Welch’s t-test, significant difference was indicated by an asterisk (* = p < 0.05). Cortisol was measured in 14 rejected and 12 controls.

Comparison of behaviours of rejected households versus controls demonstrated differences associated with stress (Table 1). We identified eight behaviours: playing, feeding, grooming, moving from one side to the other (together and apart), resting (together and apart), and head twirls. Of these, four were significantly more frequent in rejected owl monkeys, the first three are measures of whole-body stereotypies (pacing together and alone and head twirls). These stereotypic behaviours were almost not observed in control households (pacing alone p=0.0397; together p=0.0159; head twirling p=0.0476). Furthermore, feeding bouts were also more statistically frequent in rejected than controls (p=0.0238). Among other recorded behaviours, it was noted that both rejected and control juvenile owl monkeys rested together frequently and groomed very little.

**Table 1.** Total count of behaviours observed in rejected and control households. Frequency of feeding, pacing and head twirls were significantly different, being more common in rejected. Probability *p* significative to was indicated by asterisks (* = 0.05). All behaviours were scored based on camera recordings from 5 rejected and 5 control households.

| Behaviour | Rejected | Controls | *p*-value |
| --- | --- | --- | --- |
| Play | 2.00 ± 2.95 | 3.00 ± 2.17 | 0.8881 |
| Feeding | 10.00 ± 3.51 | 3.00 ± 2.30 | *0.0238 |
| Pacing alone | 11 ± 4.04 | 4 ± 3.32 | *0.0397 |
| Pacing together | 9.00 ± 6.91 | 2.00 ± 1.92 | *0.0159 |
| Resting Alone | 2.00 ± 2.77 | 1.00 ± 1.67 | 0.3095 |
| Resting together | 31.00 ± 13.95 | 17.00 ± 14.72 | 0.2222 |
| Head twirling | 6.00 ± 4.38 | 0.00 ± 0.45 | *0.0476 |
| Grooming | 0.45 ± 0.00 | 0.45 ± 0.00 | 0.9999 |

To determine the effect of maternal rejection stress on the risks of infection, we analyzed the frequency of administration of drugs to treat parasitic, bacterial and fungal infections that had been recorded in medical histories. The results are shown in Figure 2.A. Our Welch’s t-test analysis showed that the number of treatments for infections per month lived was significantly higher (p = 0.0009, two tailed, t=4.436, df=11.49) for the rejected than for the controls, who were almost never treated for infections. Furthermore, we analyzed the relationship between stress and infections with a linear regression analysis of hair cortisol concentration versus frequency of administration of drugs for infections per month lived (Figure 2.B). This showed a significant correlation (p = 0.0075), however it only explains a small portion of the variance of infection prevalence (R^2^ = 0.307, F=8.846).

**Figure 2.**
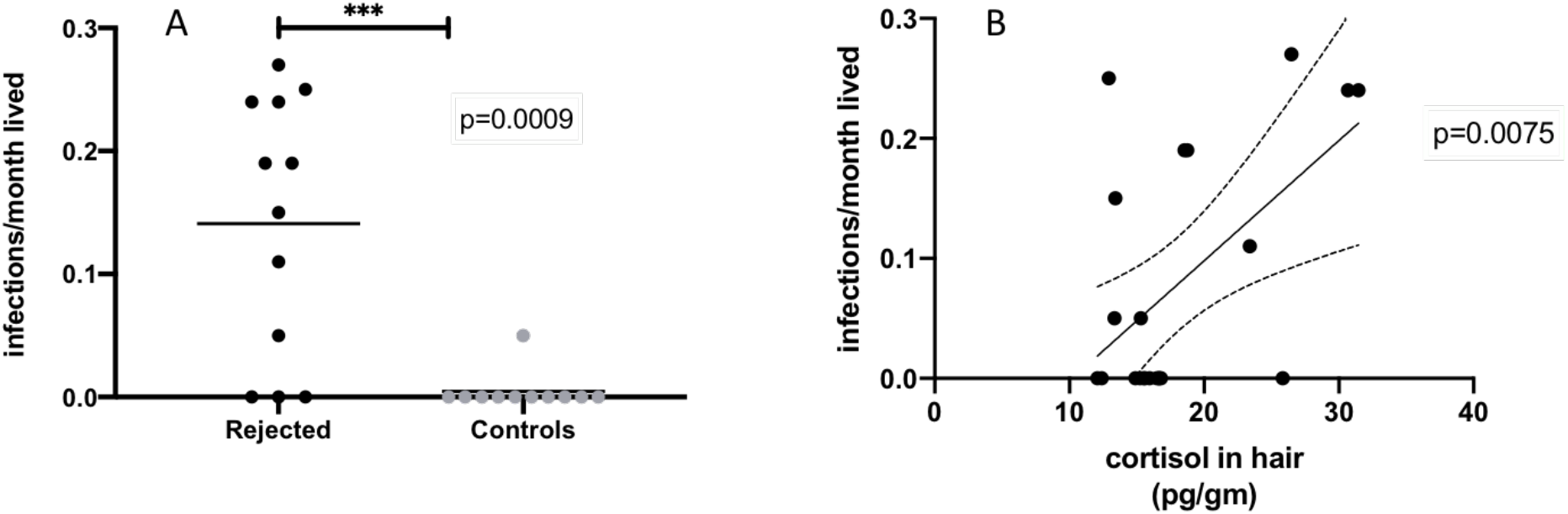
Cortisol concentration is associated with prevalence of infections in rejected juveniles but not in controls. A. Ratio of medication for infections received per month lived in rejected (in black, n=12) and control owl monkeys (in grey, n=11) B. Correlation between medication/month lived and cortisol concentration in hair across all individuals. There was a significant positive correlation between number of infection treatments per months lived and hair cortisol (linear regression p=0.0075; R^2^=0.307; F=8.846, n=23).

## 5. DISCUSSION

Our study confirmed our hypothesis that maternal rejection is associated with high stress profiles and higher frequency of infections in IVITA’s juvenile owl monkeys. Furthermore, the effects were detectable up to 2 years after exposure, even when rejected were manually raised by veterinarians. Cortisol concentrations alone did not explain the frequency of infections in this population.

Regarding the effect of stress on cortisol levels, our calculations show that the average cortisol concentration in the hair of rejected owl monkeys was 30% higher than that observed in controls (Fig.1). This suggests that in owl monkeys, maternal rejection and hand rearing leads to greater activation of the hypothalamic pituitary adrenal (HPA) axis (Sánchez 2006) despite assisted rearing of veterinarians. Our findings are consistent with experimental studies of parental separation in other species of New World monkeys, showing that circulatory cortisol increases in response to maternal deprivation in early life. For example, infant squirrel monkeys show an increase in plasma cortisol levels following temporary separation from their mothers (Levine et al., 1993). However, reports in marmosets demonstrate that changes in circulatory levels of cortisol can also occur in a different direction than ours, whereby cortisol levels in maternally separated individuals are lower than in controls (e.g. Dettling 2002a). Such variation in cortisol responses are commonly observed and could be explained by the variation in the measurement methods (cortisol in urine or blood versus hair) or in the intensity of the separation between referenced studies i.e., separation in these aforementioned studies was temporary and short-lived whereas in our study, the separation was permanent. We chose hair as a substrate to quantify circulatory cortisol, as it is considered suitable for the study of chronic stress. Cortisol is slowly deposited on the stem of the hair, therefore represents the retrospective average basal hormonal phenotype (Devenport et al., 2006; Meyer et al., 2014). Since owl monkey hair is relatively short (3-5 cm), our cortisol measurements are likely to reflect several months of slow growth. However, to confirm this finding, it is necessary to characterize the hair growth cycle for this species.

Our camera recordings confirmed that under basal conditions, stereotypy was more frequent in rejected owl monkeys (Table 1). Since both groups were raised in different environments, this is to some extent expected. The higher incidence of stereotypies suggest that emotional distress accompanies elevated and circulatory cortisol in owl monkeys. This is proposed because head twirls and pacing are classified as whole-body stereotypy (WBS), while repetitive movements are meaningless (Mason, 1991a). These behaviours have been widely associated with exposure to stressful conditions in early life, particularly when there is a disruption of the mother-child relationship (Sánchez 2006, Novak et al., 2012). In this context, recorded abnormal behaviours of rejected owl monkeys likely arise because of the experience of rejection rather than differences in breeding environments.

Rejected owl monkeys significantly fed themselves more often, which has not been reported before as a behavioural correlate to maternal deprivation. However, it has been observed under other forms of stress in primates. For example, Arce and colleagues in 2010 showed that rhesus macaques exposed to social subordination ingest more calories than dominant individuals. As intake of calorie rich food in anxious individuals can attenuate corticotropin releasing factor (CRF) in the hypothalamus, the first step in the biochemical cascade that activates the HPA axis, it has been proposed that overfeeding could reduce the effects of stress on the brain (Dallman et al., 2003). More studies are needed to confirm whether owl monkeys overfeeding behaviour is functional.

Our analysis suggests that maternal rejection is associated with the long-term frequency of infection in owl monkeys from IVITA. The weak correlation between hair cortisol and frequency of infection treatment suggests that factors other than physiological stress contribute to infection propensity in this colony. For example, Cohen et al., (2007) suggested that the stressful experience itself, modifies inflammatory and immunological responses to infections, and this is independent from cortisol secretion. Since rejected owl monkeys were fed human formula instead of breast milk, and were hand reared by humans, it is possible they were more susceptible to develop infections and were more exposed to pathogens than controls.

However, studies have shown that infection rates in primates are increased when they are simply exposed to poor quality maternal care, even when they are not hand reared by humans (Kinnally et al., 2019). Therefore, it is likely that elevated infection rates in spontaneously rejected owl monkeys are the result of a combination of factors which show plasticity to maternal stress. Principal candidates to mediate this phenomenon are epigenetic changes in genes involved in stress reactivity (i.e. serotonin transporter gene) which can predict differences in adulthood infection rates in rhesus macaques exposed to maternal stress in early life (Kinnally; 2014).

In conclusion, maternal rejection is a strong developmental stressor for IVITA owl monkeys. Given our findings, considering the individual history of rejection and monitoring stress levels could help veterinarians to identify monkeys most likely to develop clinical symptoms of infection and improve the management of this species in captivity.

